# TMEM106B, a risk factor for FTLD and aging, has an intrinsically disordered cytoplasmic domain

**DOI:** 10.1101/317958

**Authors:** Jian Kang, Liangzhong Lim, Jianxing Song

## Abstract

TMEM106B was initially identified as a risk factor for FTLD, but recent studies highlighted its general role in neurodegenerative diseases. Very recently TMEM106B has also been characterized to regulate aging phenotypes. TMEM106B is a 274-residue lysosomal protein whose cytoplasmic domain functions in the endosomal/autophagy pathway by dynamically and transiently interacting with diverse categories of proteins but the underlying structural basis remains completely unknown. Here we conducted bioinformatics analysis and biophysical characterization by CD and NMR spectroscopy, and obtained results reveal that the TMEM106B cytoplasmic domain is intrinsically disordered with no well-defined three-dimensional structure. Nevertheless, detailed analysis of various multi-dimensional NMR spectra allowed defining residue-specific conformations and dynamics. Overall, the TMEM106B cytoplasmic domain is lacking of any tight tertiary packing and relatively flexible. However, several segments are populated with dynamic/nascent secondary structures and have relatively restricted backbone motions. In particular, the fragment Ser12-Met36 is highly populated with α- helix conformation. Our study thus decodes that being intrinsically disordered allows the TMEM106B cytoplasmic domain to dynamically and transiently interact with a variety of distinct partners.

## Introduction

Transmembrane protein 106B (TMEM106B) is 274-residue protein whose gene locus was initially identified by a genome-wide association study to be a risk factor for the occurrence of frontotemporal lobar degeneration (FTLD), which is the second most common form of progressive dementia in people under 65 years (1,2). In 2012 it was reported that TMEM106B SNPs might be also critical for the pathological presentation of Alzheimer disease (3). Remarkably, the risk allele of TMEM106B was found to accompany significantly reduced volume of the superior temporal gyrus, most markedly in the left hemisphere (4), which includes structures critical for language processing, usually affected in FTLD patients (5). TMEM106B was also reported to be a genetic modifier of disease in the carriers of C9ORF72 expansions (6). Very recently, differential aging analysis further identified that variants of TMEM106B are responsible for aging phenotypes. Briefly, TMEM106B risk variants were shown to be associated with inflammation, neuronal loss, and cognitive deficits for older individuals (> 65 years) even without any known brain disease, and their impact is particularly selective for the frontal cerebral cortex (7).

TMEM106B is a member of the TMEM106 family of proteins of 200-300 amino acids with largely unknown function, which consists of TMEM106A, TMEM106B, and TMEM106C. Immunohistochemical assessments showed that TMEM106B is expressed in neurons as well as in glial and endothelial cells (8). The expression levels of MEM106B appear to be critical for lysosome size, acidification, function, and transport in various cell types; and its upregulation leads to lysosomal defects such as improper lysosomal formation, poor lysosomal acidification, reduced lysosomal transport, and increased lysosomal stress (9–14). These studies revealing the functions of neuronal TMEM106B in regulating lysosomal size, motility and responsiveness to stress, therefore underscore the key role of lysosomal biology in FTLD and aging.

Recently, biochemical characterizations revealed that TMEM106B is a single-pass, type 2 integral membrane glycoprotein predominantly located in the membranes of endosomes and lysosomes with a transmembrane domain predicted to be over residues 96– 118 (Fig. 1A). However, the molecular mechanisms underlying the physiological functions of MEM106B remain largely elusive. So far, no protein binding partner has been identified to interact with the luminal C-terminus of TMEM106B. On the other hand, the N-terminal cytoplasmic domain of TMEM106B has been amazingly implicated in dynamically and transiently interacting with very diverse proteins or complexes, which include endocytic adaptor proteins such as clathrin heavy chain (CLTC) (12); proteins including CHMP2B of the endosomal sorting complexes required for transport III (ESCRT-III) complex (13); and microtubule-associated protein 6 (Map6) (14). In this regard, characterization of its high-resolution structure is expected to provide a key foundation for addressing how the TMEM106B cytoplasmic domain can interact with significantly distinctive partners.

**Figure 1.**
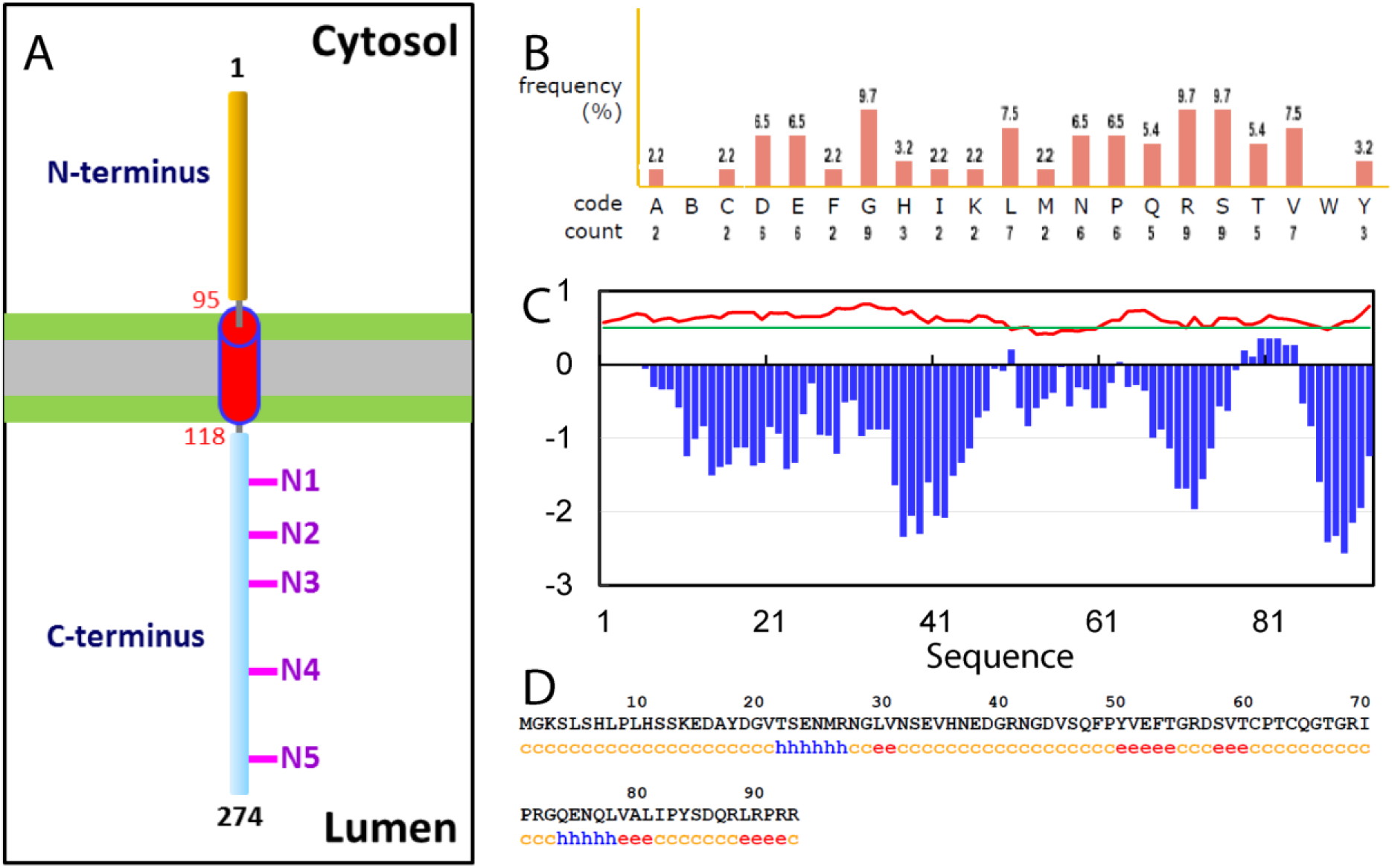
Bioinformatics analysis of the TMEM106B cytoplasmic domain. (A) A model of the TMEM106B transmembrane topology. N1–N5 indicates the five potential glycosylation motifs within the luminal C-terminus of TMEM106B. (B) Amino acid composition of the TMEM106B cytoplasmic domain. (C) Kyte & Doolittle Hydrophobicity Score (blue bars); and Disorder Score (red line) predicted by IUPred program, which ranges between 0 and 1. Scores above 0.5 (green line) indicate disorder. (D) Secondary structures predicted by GOR4 program: c: random coil, h: helix and e: extended strand.

So far no biophysical/structural study has been reported on the cytoplasmic domain of TMEM106B in the free state, or in complex with binding partners. In the present study, we aimed to determine its solution conformation and dynamics by CD and NMR spectroscopy. Our results reveal that despite having relatively random sequence, the TMEM106B cytoplasmic domain is intrinsically disordered without any stable secondary and tertiary structures. Nevertheless, residue-specific NMR probes allowed the identification of several regions of the TMEM106B cytoplasmic domain which are weakly populated with either helical or extended conformations to different extents. Therefore, our present study deciphers that being intrinsically disordered appears to offer a unique capacity for the TMEM106B cytoplasmic domain to interact with diverse binding partners dynamically and transiently, which appears to be required for implementing its physiological functions.

## Materials and Methods

### Bioinformatics analysis, cloning, expression and purification of the cytoplasmic domain of TMEM106B

The sequence of the TMEM106B cytoplasmic domain over residues 1-93 was extensively analyzed by various bioinformatics programs to assess its hydrophobicity (15), Disorder score by IUPred (16), and secondary structures by GOR4 (17).

The DNA encoding residues 1-93 of TMEM106B was purchased from GeneScript and subsequently cloned into a modified vector pET28a with a C-terminal His-tag (18,19). The recombinant protein was found in supernatant and thus first purified by Ni^2+^-affinity column under native condition followed by further purification with FPLC on a Superdex-200 column. To produce isotope-labeled proteins for NMR studies, the same procedures were used except that the bacteria were grown in M9 medium with addition of (^15^NH_4_)_2_SO_4_ for ^15^N-labeling; and (^15^NH_4_)_2_SO_4_/^13^C-glucose for ^15^N-/^13^C-double labeling (18,19).

### CD and NMR experiments

All CD experiments were performed on a Jasco J-1500 spectropolarimeter equipped with a thermal controller (18,19). Far-UV CD spectrum was collected on a protein sample at 10 μM in 5 mM phosphate buffer (pH 6.5) using 1-mm path length cuvettes, while near-UV CD spectra in the absence and presence of 8 M urea were acquired on protein samples at 100 μM in the same buffer with 10-mm path length cuvettes. Data from five independent scans were added and averaged. Deconvolution of the far-UV CD spectrum to estimate the contents of secondary structures was conducted by K2D2 program (20).

All NMR experiments were acquired on an 800 MHz Bruker Avance spectrometer equipped with pulse field gradient units as described previously (18,19). NMR experiments including HNCACB and CCC(CO)NH were acquired on ^15^N-/^13^C-double labelled samples for achieving sequential assignments, while ^15^N-edited HSQC-TOCSY and HSQC-NOESY were collected on ^15^N-labelled samples to identify NOE connectivities at protein concentrations of 200 μM in 5 mM phosphate buffer (pH 6.5). For assessing the backbone dynamics on the ps-ns time scale, {^1^H}-^15^N steady-state NOEs were obtained by recording spectra on ^15^N-labeled samples at 200 μM in 5 mM phosphate buffer (pH 6.5) as previously described (18,19). Obtained chemical shifts were further analysed by SSP program (21) to obtain residue-specific score of secondary structure propensity (SSP).

## Results

### Bioinformatics analysis of the TMEM106B cytoplasmic domain

In the present study, to facilitate experimental investigations, the cytoplasmic domain of TMEM106B was first assessed by a variety of bioinformatics tools to gain insights into its amino acid composition, ordered/disordered regions and secondary structure. As shown in Fig. 1B, it has a relatively random composition of amino acids similar to the TDP-43 N-terminal domain adopting an ubiquitin-like fold (18), which is not highly biased to a certain type of amino acids as observed in the TDP-43 prion-like domain (19). On the other hand, except for residues Tyr50 and Leu78-Tyr84 with small positive hydrophobicity scores, all other residues have negative hydrophobicity scores (Fig. 1C), suggesting that it is overall hydrophilic. Furthermore, the disorder scores of most residues predicted by IUPred are > 0.5 (Fig. 1C), implying that it might be intrinsically disordered (16,22). Indeed, based on the secondary structure prediction result (Fig. 1D), a large portion of the sequence is in random coil conformation. However, two very short fragments, namely Thr22-Arg27 and Gln74- Leu78, are predicted to form α-helices, while five short fragments, namely Leu30-Val31, Tyr50-Thr54, Ser58-Thr60, Val79-Leu81 and Leu89-Pro91, adopt extended conformations.

### Conformational properties of the TMEM106B cytoplasmic domain

The recombinant protein of the TMEM106B cytoplasmic domain is highly soluble in buffer at neutral pH. Its far-UV CD spectrum has the negative signal maximum at ~208 nm and a shoulder negative signal at ~222 nm, as well as a positive signal at ~190 nm, which indicates that it contains helical conformations to some degree. We thus conducted further deconvolution of the CD spectrum by K2D2 program (20), which estimated the TMEM106B cytoplasmic domain to contain 35.7% α-helix, 13.2% β-stands and 51.1% random coil.

While far-UV CD spectroscopy reflects secondary structure features of a protein, near-UV CD is very sensitive to the tight tertiary packing (22). Therefore, we subsequently recorded the near-UV CD spectra of the TMEM106B cytoplasmic domain in the absence and presence of 8 M denaturant urea (Fig. 2B). The high similarity between two near-UV spectra clearly suggested that it is lacking of any tight tertiary packing even under the native condition (22). Completely consistent with this results, its two-dimensional ^1^H-^15^N NMR HSQC spectrum has very narrow spectral dispersions over both dimensions, with only ~0.9 ppm over ^1^H and ~19 ppm over ^15^N dimensions (Fig. 2C). Therefore, preliminary CD and NMR characterizations revealed that the TMEM106B cytoplasmic domain is an intrinsically disordered domain (IDD), which is lacking of any tight tertiary packing but populated with secondary structures to different extents.

**Figure 2.**
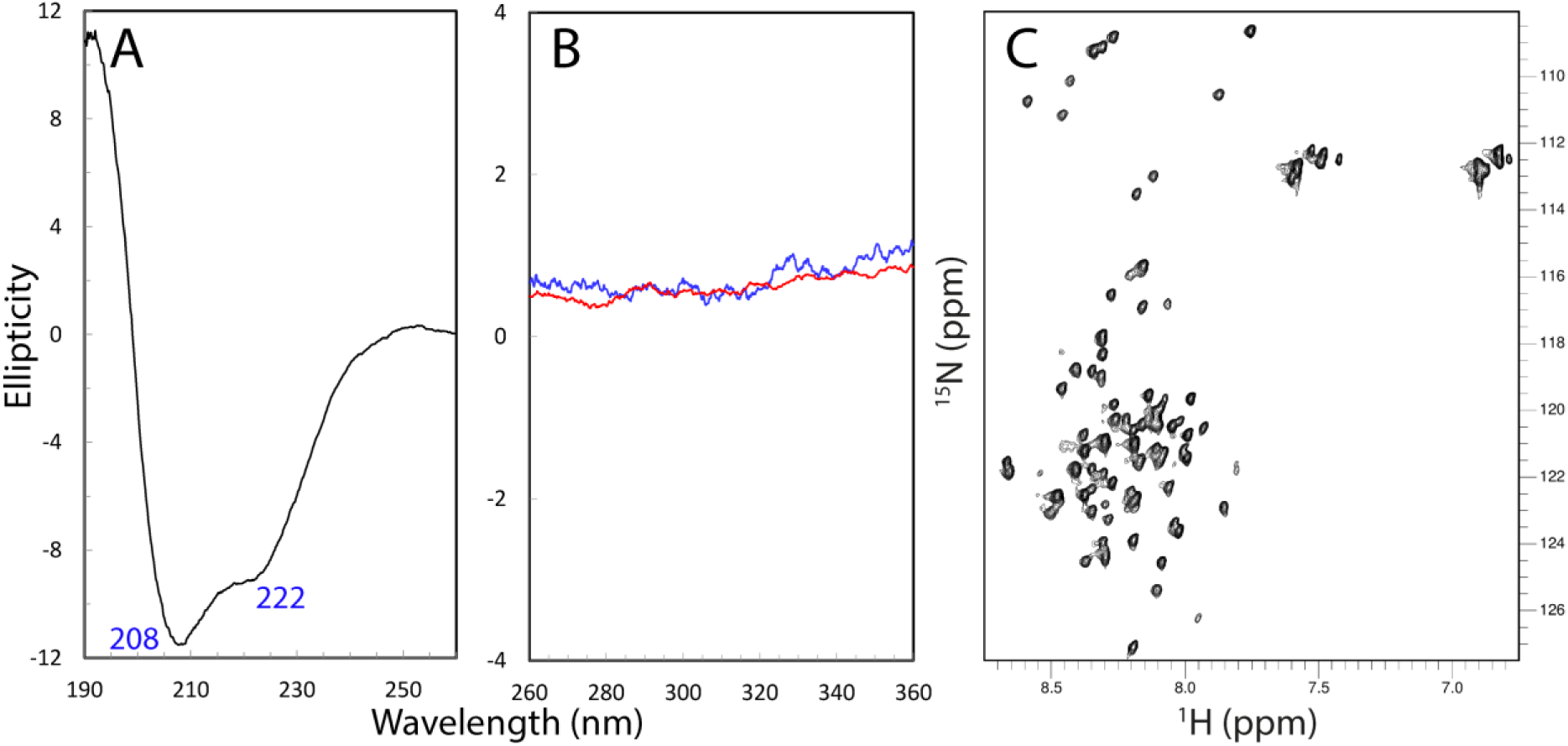
Biophysical characterization of the TMEM106B cytoplasmic domain. (A) Far-UV CD spectrum of the TMEM106B cytoplasmic domain at a protein concentration of 10 μM at 25 °C in 5 mM phosphate buffer (pH 6.5). (B) Near-UV CD spectra of the TMEM106B cytoplasmic domain at a protein concentration of 100 μM at 25 °C in the 5 mM phosphate buffer (pH 6.5) in the absence (blue) and in the presence of 8 M urea. (C) Two-dimensional ^1^H-^15^N NMR HSQC spectrum of the TMEM106B cytoplasmic domain acquired at 25 °C at a concentration of 100 μM in 5 mM phosphate buffer (pH 6.5).

### Residue-specific conformation and dynamics of the TMEM106B cytoplasmic domain

NMR spectroscopy is the most powerful biophysical tool to pinpoint residual structures existing in the unfolded proteins (18,19,22,23). Therefore, in order to gain insights into residue-specific conformational of the TMEM106B cytoplasmic domain, we acquired and analyzed three-dimensional NMR spectra including CCC(CO)NH, HN(CO)CACB, HSQC-TOCSY and HSQC-NOESY spectra. Subsequently we have successfully achieved sequential assignments of all residues except for the first residue and six Pro residues without HSQC peaks, whose side chain resonances, however, were also assigned. Fig. 3A presents (ΔCα-ΔCβ) chemical shift, which is a sensitive indicator of the residual secondary structures in disordered proteins (18,19, 23). The very small absolute values of (ΔCα-ΔCβ) over the whole sequence indicate that the TMEM106B cytoplasmic domain is lacking of well-formed secondary structures. Nevertheless, several regions show relatively large deviations, thus suggesting that these regions may be populated with dynamic secondary structures to some degree (Fig. 2A).

**Figure 3.**
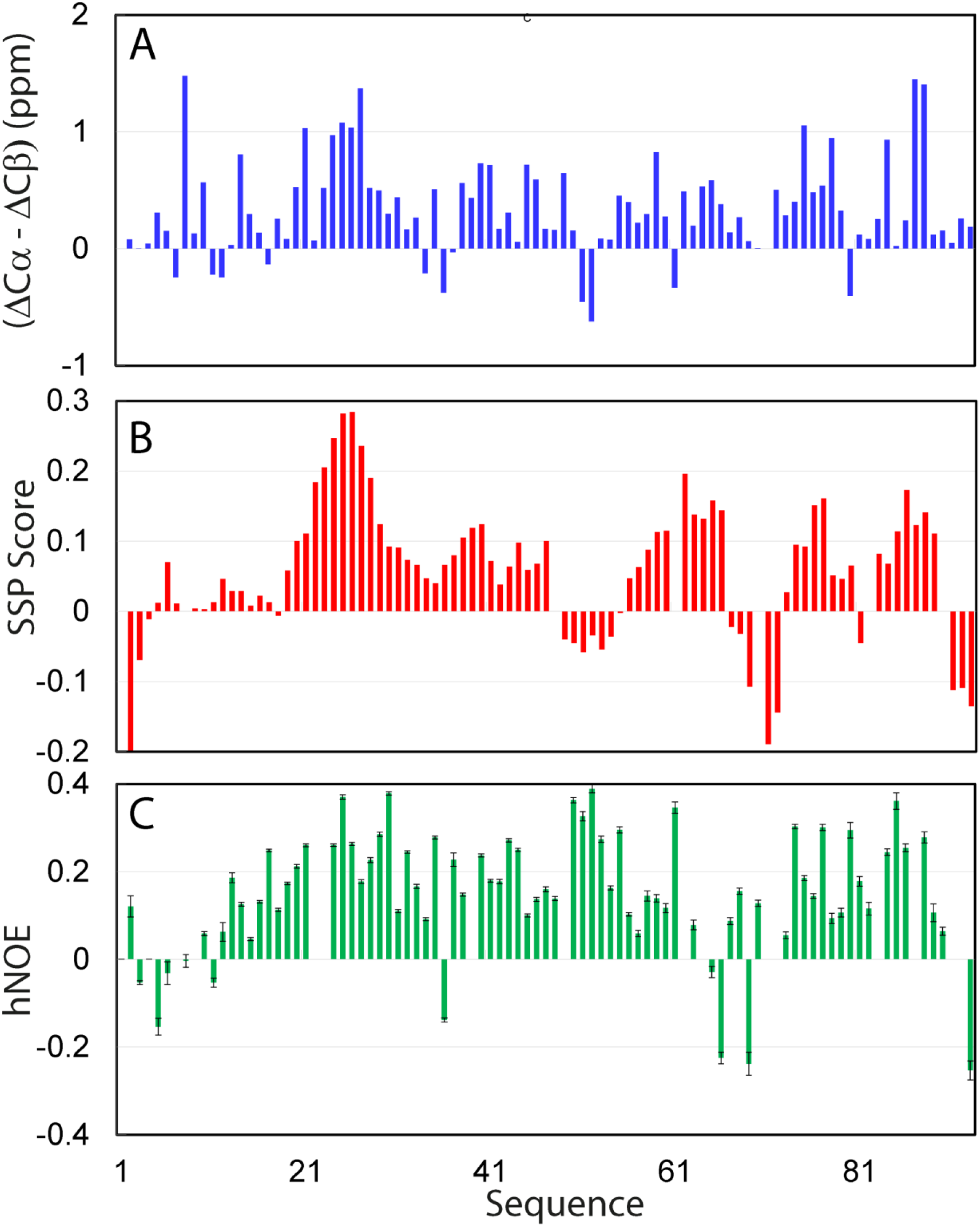
Residue-specific conformation of the TMEM106B cytoplasmic domain. (A) Residue specific (ΔCα-ΔCβ) chemical shifts of the TMEM106B cytoplasmic domain. (B) Secondary structure propensity (SSP) score obtained by analyzing chemical shifts of the TMEM106B cytoplasmic domain with the SSP program. A score of +1 is for the well-formed helix while a score of -1 for the well-formed extended strand. (C) {^1^H}-^15^N heteronuclear steady-state NOE (hNOE) of the TMEM106B cytoplasmic domain.

To obtain quantitative insights into the populations of different secondary structures, we further analyzed chemical shifts of the TMEM106B cytoplasmic domain by SSP program (21), and the results are shown in Fig 3B. Indeed, all residues have the absolute values of SSP scores less than 0.3, confirming that the whole domain has no well-formed secondary structure. Nevertheless, several regions including Gly20-Gly29, Glu38-Gly40, Val59-Gly66, Asn76-Gln77 and Ser85-Leu89 have residue with SSP scores larger than 0.1, thus implying that they are populated with dynamic helical conformations. On the other hand, several very short segments including Gly3, Arg69-Arg72 and Pro91-Arg93 also have SSP scores < -0.1, implying that these regions might have intrinsic capacity to adopt extended conformations.

We also assessed the backbone rigidity of the TMEM106B cytoplasmic domain by collecting heteronuclear NOEs, which provides a measure to the backbone flexibility on the ps-ns timescale (18,19,23). As shown in Fig 3C, the backbone appears to be overall flexible on ps-ns time scale as judged from the relatively small or even negative heteronuclear NOEs (hNOEs), with an average value of only 0.15 (Fig. 3C), which is much smaller than that for a well-folded protein (18). Nevertheless, this average value is much larger than those for the unfolded states of ALS-causing SOD1 with an average value of -0.1 (23) and C71G-PFN1 with an average value of -0.15 (24) also collected at 800 MHz. Therefore, it appears that the backbone motions on ps-ns time scale are partially restricted most likely due to the formation of dynamically-populated secondary structures, in particular helical conformations over several fragments.

Indeed, further analysis of HSQC-NOESY spectrum allowed the identification of many sequential and medium-range NOE connectivities (NOEs) characteristic of helical conformations, which include dNN(i, i+1), dαN(i, i+2), dNN(i, i+2), dαN(i, i+3) and even dαN(i, i+4) NOEs (Fig. 4). Briefly, generally consistent with SSP prediction (Fig. 3B), the long fragment over Ser12-Asp57 appears to be populated with helical conformations. In particular, the fragment Ser12-Met36 is highly populated with α-helical conformations as judged from the existence of many NOEs characteristic of α-helical conformation, which even has many dNN(i, i+2), dαN(i, i+3) and dαN(i, i+4) NOEs. It is also interesting to note that several other fragments also have dαN(i, i+2) NOEs, implying that these regions may be populated with dynamic helical/loop conformations. On the other hand, no long-range NOE was found, suggesting that the TMEM106B cytoplasmic domain is lacking of any tight tertiary packing.

**Figure 4.**
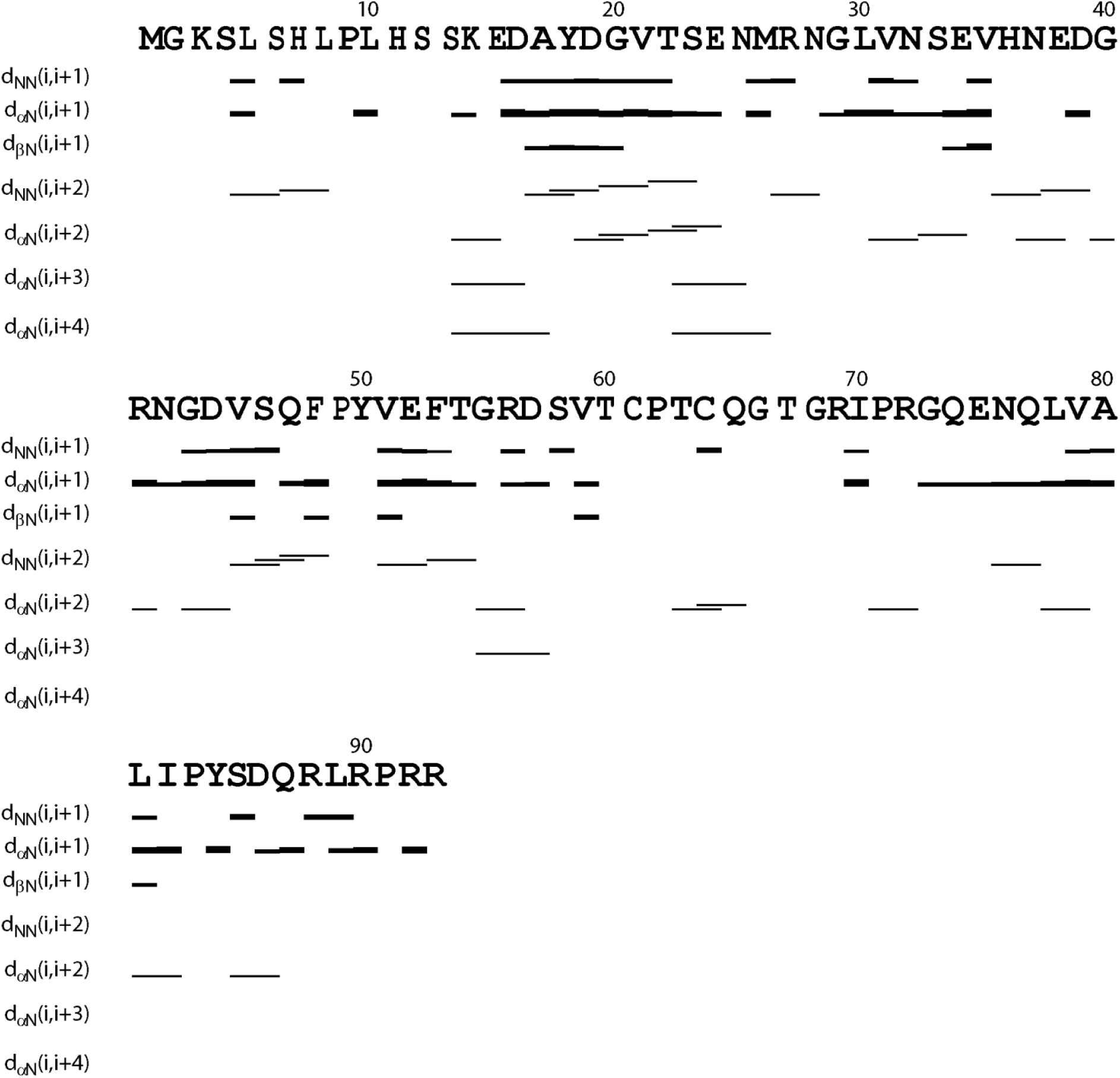
Characteristic NOE connectivities defining secondary structures of the TMEM106B cytoplasmic domain.

## Discussion

Although TMEM106B variants were initially identified as risk factors for FTLD (1), recent studies suggest that it may have more general involvements in neurodegenerative diseases including Alzheimer disease (3), and other in carriers of C9ORF72 expansions such as amyotrophic lateral sclerosis (ALS) (6). Intriguingly, TMEM106B variants have been very recently identified to regulate aging phenotypes, thus implying that the normal aging and neurodegenerative disorders might have some convergent mechanisms/pathways (7). Indeed, TMEM106B has been characterized to be a lysosomal protein whose variants are associated with defects of the endolysosomal pathway, supporting the emerging view that lysosomal biology plays key roles in neurodegenerative diseases and aging (1–14).

The TMEM106B cytoplasmic domain functions in the endosomal/autophagy pathway by interacting with various protein partners. Briefly, the TMEM106B cytoplasmic domain has been functionally mapped out to interact with three diverse categories of proteins: endocytic adaptor proteins (12); proteins of the endosomal sorting complexes (13); and microtubule-associated protein 6 (Map6) (14). One emerging feature characteristic of these interactions is their dynamic and transient nature. In fact, the risk factor T185 was shown to be associated with CHMP2B more tightly than S185, thus slightly reducing autophagic flux (13). Strikingly, the TMEM106B cytoplasmic domain appears to bind the intrinsically disordered region of MAP6 (14).

Elucidation of the three-dimensional structure of the TMEM106B cytoplasmic domain will not only provide mechanistic insights how the small TMEM106B cytoplasmic domain can interact with such diverse partners, but may also offer the clues for future discovery/design of molecules to interfere in the interactions for therapeutic applications. In the present study, we first assessed the structural properties of the TMEM106B cytoplasmic domain by both bioinformatics analysis and biophysical characterization including CD and NMR HSQC spectroscopy. The results together indicated that the TMEM106B cytoplasmic domain is an intrinsically disordered domain with no well-folded three-dimensional structure.

Nevertheless, by analyzing a set of multi-dimensional NMR spectra, we have successfully obtained residue-specific NMR parameters, which thus define conformational and dynamic properties of the TMEM106B cytoplasmic domain. Overall, the TMEM106B cytoplasmic domain is lacking of any tight tertiary packing as evidenced by the lack of any long-range NOE, as well as relatively flexible as judged by its small average value of hNOEs. However, many segments are populated with dynamic/nascent secondary structures and have relatively restricted backbone motions. For example, a long region over Ser12-Asp57 is populated with helical conformations and particularly the fragment Ser12-Met36 is highly populated with α-helical conformation. Additionally several other fragments are also populated with dynamic helical/loop conformations.

Our present study thus offers the structural basis for the TMEM106B cytoplasmic domain to dynamically and transiently interact with such diverse protein partners. It has been well-established that one unique advantage for a protein to be intrinsically disordered is the capacity in interacting with a large and diverse set of binding partners dynamically and transiently (26). One scenario is “binding-induced folding”: the intrinsically disordered domain with dynamically populated secondary structures becomes folded only upon binding to its partners, as we previously found on the regulatory domain of PAK4 kinase, which is intrinsically disordered but forms a well-defined helix in complex with the kinase domain (27). Alternatively, the intrinsically disordered domain may still remain largely disordered and dynamic even upon forming a “fuzzy complex” with its partners, as we previously observed on the hepatitis C virus NS5A protein which could form a “fuzzy complex” with the human host factor VAPB (27). Therefore, upon binding to its partners the TMEM106B cytoplasmic domain might fold into a well-defined three-dimensional structure, or still remain disordered and dynamic. However, it is also possible that the TMEM106B cytoplasmic domain folds into a well-defined structure upon binding to one or one category of partners, while remain largely disordered and dynamic after interacting with another or another category of partners. Therefore in the future, it is of fundamental and therapeutic interest to devote extensive efforts to first mapping out the binding domains/regions of the partners of the TMEM106B cytoplasmic domain, and to subsequently determining their complex structures.

## Acknowledgement

This study is supported by Ministry of Education of Singapore (MOE) Tier 2 Grant MOE2015-T2-1-111 to Jianxing Song.

## References

1. Van Deerlin V.M., Sleiman P.M., Martinez-Lage M., et al., Common variants at 7p21 are associated with frontotemporal lobar degeneration with TDP-43 inclusions. Nat Genet 42 2010. 234–239.

2. Nicholson A.M., Rademakers R. What we know about TMEM106B in neurodegeneration. Acta Neuropathol. 132 2016. 639–651.

3. Rutherford N.J., Carrasquillo M.M., Li M., et al., TMEM106B risk variant is implicated in the pathologic presentation of Alzheimer disease. Neurology 79 2012. 717–718.

4. Adams H.H., Verhaaren B.F., Vrooman H.A., et al., TMEM106B influences volume of left-sided temporal lobe and interhemispheric structures in the general population. Biol Psychiatry 76 2014. 503–508.

5. Mackenzie I.R., Neumann M., Baborie A., et al., A harmonized classification system for FTLD-TDP pathology. Acta Neuropathol 122 2011. 111–113.

6. Gallagher M.D., Suh E., Grossman M., et al., TMEM106B is a genetic modifier of frontotemporal lobar degeneration with C9orf72 hexanucleotide repeat expansions. Acta Neuropathol. 127 2014. 407–18.

7. Rhinn H., Abeliovich A. Differential Aging Analysis in Human Cerebral Cortex Identifies Variants in TMEM106B and GRN that Regulate Aging Phenotypes. Cell Syst. 4 2017. 404–415.e5.

8. Nicholson A.M., Finch N.A., Wojtas A., et al., TMEM106B p.T185S regulates TMEM106B protein levels: implications for frontotemporal dementia. J Neurochem. 126 2013. 781–791.

9. Chen-Plotkin A.S., Unger T.L., Gallagher M.D., et al., TMEM106B, the risk gene for frontotemporal dementia, is regulated by the micro-RNA-132/212 cluster and affects progranulin pathways. J Neurosci. 32 2012. 11213–11227

10. Lang C.M., Fellerer K., Schwenk B.M., et al., Membrane orientation and subcellular localization of transmembrane protein 106B TMEM106B., a major risk factor for frontotemporal lobar degeneration. J Biol Chem. 287 2012. 19355–19365.

11. Brady O.A., Zheng Y., Murphy K., et al., The frontotemporal lobar degeneration risk factor, TMEM106B, regulates lysosomal morphology and function. Hum Mol Genet 22 2012. 685–695.

12. Stagi M., Klein Z.A., Gould T.J., et al., Lysosome size, motility and stress response regulated by fronto-temporal dementia modifier TMEM106B. Mol Cell Neurosci 61 2014. 226–240.

13. Jun M.H., Han J.H., Lee Y.K., et al., TMEM106B, a frontotemporal lobar dementia FTLD. modifier, associates with FTD-3-linked CHMP2B, a complex of ESCRTIII. Mol Brain 8 2015. 85.

14. Schwenk B.M., Lang C.M., Hogl S., et al., The FTLD risk factor TMEM106B and MAP6 control dendritic trafficking of lysosomes. EMBO J. 33 2014. 450–467.

15. Kyte J., Doolittle R.F. A simple method for displaying the hydropathic character of a protein. J Mol Biol 157 1982. 105–132.

16. Dosztányi Z., Csizmók V., Tompa P., et al., IUPred: web server for the prediction of intrinsically unstructured regions of proteins based on estimated energy content. Bioinformatics 21 2005. 3433–3434.

17. Garnier J., Gibrat J.F., Robson B. GOR secondary structure prediction method version IV. Methods Enzymol. 266 1996. 540–553.

18. Qin H., Lim L.Z., Wei Y., et al. TDP-43 N terminus encodes a novel ubiquitin-like fold and its unfolded form in equilibrium that can be shifted by binding to ssDNA. Proc. Natl. Acad. Sci. U. S. A., 111 2014., pp. 18619–18624.

19. Lim L., Wei Y., Lu Y., et al. ALS-causing mutations significantly perturb the self-assembly and interaction with nucleic acid of the intrinsically disordered prion-like domain of TDP-43. PLoS Biol., 14 2016. e1002338

20. Perez-Iratxeta C., Andrade-Navarro M.A. K2D2: estimation of protein secondary structure from circular dichroism spectra. BMC Struct Biol. 8 2008. 25.

21. Marsh J.A., Singh V.K., Jia Z., et al. Sensitivity of secondary structure propensities to sequence differences between alpha- and gamma-synuclein: implications for fibrillation. Protein Sci. 15 2006. 2795–2804.

22. Li M., Song J. The N- and C-termini of the human Nogo molecules are intrinsically unstructured: bioinformatics, CD, NMR characterization, and functional implications. Proteins. 68 2007. 100–8.

23. Dyson H.J., Wright P.E. Unfolded proteins and protein folding studied by NMR. Chem. Rev. 104 2004. 3607–3622.

24. Lim L., Song J. SALS-linked WT-SOD1 adopts a highly similar helical conformation as FALS-causing L126Z-SOD1 in a membrane environment. Biochim Biophys Acta. 1858 2016. 2223–2230.

25. Lim L., Kang J., Song J. ALS-causing profilin-1-mutant forms a non-native helical structure in membrane environments. Biochim Biophys Acta. 1859 2017. 2161–2170.

26. Wright P.E., Dyson H.J. Intrinsically disordered proteins in cellular signalling and regulation. Nat Rev Mol Cell Biol. 16 2015. 18–29.

27. Wang W., Lim L., Baskaran Y., et al. NMR binding and crystal structure reveal that intrinsically-unstructured regulatory domain auto-inhibits PAK4 by a mechanism different from that of PAK1. Biochem Biophys Res Commun. 438 2013. 169–174.

28. Gupta G., Qin H., Song J. Intrinsically unstructured domain 3 of hepatitis C Virus NS5A forms a “fuzzy complex” with VAPB-MSP domain which carries ALS-causing mutations PloS ONE 7 6. e39261 2012.

